# Soft X-ray tomography reveals variations in *B.subtilis* biofilm structure upon *tasA* deletion

**DOI:** 10.1101/2024.10.01.615898

**Authors:** Anthoula Chatzimpinou, Anne Diehl, A. Tobias Harhoff, Kristina Driller, Bieke Vanslembrouck, Jian-Hua Chen, Kristaps Kairiss, Valentina Loconte, Mark A Le Gros, Carolyn Larabell, Kürşad Turgay, Hartmut Oschkinat, Venera Weinhardt

## Abstract

Bacterial biofilms are complex communities of cells within a self-produced extracellular matrix. They play crucial roles in healthcare, nutrition, agriculture and environmental research, yet an analysis of their elaborate 3D architecture remains challenging. Understanding mechanisms of biofilm formation, particularly the effects of chemical, physical, and genetic influences or modifications, is crucial but requires structural information at subcellular resolution to enable a community-level analysis of biofilms. In this work, we developed a “biofilm-in-capillary” growth method compatible with full-rotation soft X-ray tomography, providing high-resolution 3D imaging of bacterial cells and their surrounding extracellular matrix during biofilm formation, without drying or fixation steps. This approach offers 50 nm isotropic spatial resolution, rapid imaging time, and quantitative native analysis of biofilm structure. We demonstrate the potential of our method using *Bacillus subtilis* biofilms, detecting coherent alignment and chaining of wild-type cells while they are travelling towards the oxygen-rich capillary tip region. In stark contrast, the genetic knock-out Δ*tasA* shows a loss of cellular orientation, including changes in the extracellular matrix in volume and chemical density. Notably, we show that the addition of TasA protein to a culture of a Δ*tasA* strain restores the extracellular matrix density and leads to a compaction of cell assemblies, yet no chaining is observed as for the wildtype. Our approach to imaging biofilms is scalable and transferable, opening new avenues for examining biofilm structure and function across various species, including mixed biofilms, and observing 3D reorganization in response to genetic and environmental factors.

## INTRODUCTION

Bacterial biofilms are constituted by various macromolecules such as exopolysaccharides (EPS), DNA, and a major protein component that forms filaments or fibrils^1^. This heterogeneous mixture is complemented by small and medium-sized compounds, among them nutrients, signaling molecules, and surfactants^2^. Although biofilms in natural environments are inhabited by a variety of bacteria and other organisms, structural investigations on model biofilms, e.g. of *Bacillus subtilis*, help to understand basic principles of construction and function. *B. subtilis* produces different types of biofilms depending on culture conditions, such as submerged biofilm within a liquid or pellicles that are formed on surfaces of liquids^3^. They have been extensively studied with respect to extracellular matrix (ECM) composition and function^4^ and structural properties of the biofilm proteins TasA^5,6^ and TapA^7^. The major proteinaceous biofilm component TasA can form different oligomeric species, including filaments, by a strand complementation mechanism^5,7^.

Different methodological attempts have been made to analyze the entire biofilm architecture using mass spectrometry^8^, magnetic resonance imaging^9,10^, scanning transmission X-ray microscopy^11,12^, small and wide-angle X-ray scattering^13^ or different electron microscopic techniques^14^. Biofilms are heterogeneous assemblies containing up to 97 % (w/w) water^15^, hence all fixation, drying, freezing and dehydration steps compromise morphology and 3D structure, representing a challenge for 3D imaging techniques. For example, ultrastructural characterization of biofilm matrix and embedded bacterial cells with scanning electron microscopy requires additionally to special and careful protocols 3D reconstruction tools^14^. So far, there are no reports on high-resolution imaging of biofilms with techniques that did not require sample modification.

Here, we employ soft X-ray tomography (SXT) as a label-free imaging modality with a spatial resolution of 25-60 nm to investigate the role of the essential biofilm protein TasA. To understand how *tasA* deletion affects the biofilm 3D architecture, we developed a biofilm-in-capillary workflow using *B. subtilis* WT and *ΔtasA* strains as examples. By 3D imaging, we investigate i) the collective patterning of bacterial cells in biofilms, ii) changes in ECM distribution, and iii) phenotypical changes of individual bacteria in suspension.

During the process of data acquisition, a set of X-ray projection images (shadows) are collected at different rotation angles around a cylindrical sample represented by a very narrow capillary (typical tip diameter 10-12 μm). Each data acquisition is followed by 3D reconstruction which generates a volume of a specimen with a resolution that may approach 25-60 nm.

SXT operates in the “water window” of the electromagnetic spectrum, exploiting the range between carbon and oxygen absorption edges (4.4 nm to 2.3 nm wavelength) for natural contrast of carbon-rich materials and transparency of oxygen-rich aqueous media^16,17^. In this energy range, photoelectric absorption is the most dominant process, such that the concentration of chemical species relates to the attenuation of X-rays by the Beer-Lambert law, *I*(*z*) *= I*_0_*e*^−µ*z*^, where *µ* is the linear absorption coefficient (or shortly LAC) of the material with thickness *z*. Scattering in the “water window” energy range is negligible hence the LAC is approximated as 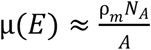, where ρ_*m*_, *N*_*A*_ and *A* are the mass density, Avogadro’s number and the atomic mass number, respectively^18^. As a constant X-ray energy of 530 eV is used, the LAC measurement is a function of molecular composition (atomic number and mass) and concentration (mass density). This unique quantitative nature of SXT has been employed to distinguish cellular organelles^19–21^, to detect variations in DNA packing^22^, and to study states of viral replication^23^. As an advantage, SXT does not require labelling, fixation or staining, enabling the unperturbed investigation of samples that contain a high amount of water.

In this work, we develop a “biofilm-in-capillary” growth method compatible with full-rotation SXT of up to 200 μm-thick biofilms with a spatial resolution of 50 nm. Using a machine learning approach, we incorporate single-cell and ECM segmentation for systematic analysis of individual cell phenotypes, their spatial organization, and density of ECM. Employing SXT, we were able to detect and analyze the loss of cell orientation after deletion of *tasA*. Upon rescue with TasA protein, we could detect partial restoration of the ECM structure. We explore correlated changes in ECM structure and cell phenotypes and discuss the role of TasA in biofilm formation. Altogether, we show that the developed biofilm-in-capillary full-rotation SXT workflow is an efficient method for the visualization and analysis of biofilms at the subcellular level under varying conditions.

## RESULTS

### Biofilm-in-capillary workflow, applied to B. subtilis

To optimize the experimental conditions for full rotation SXT, we developed a workflow that allows *B. subtilis* to form biofilms inside thin-walled glass capillaries (Figure 1a), while ensuring that oxygen and medium optimal for lipopeptide production (MOLP) are provided^24,25^. Starting from a standard *B. subtilis* culture in Luria-Bertani (LB) medium, we inoculated a 6-well plate with the bacteria diluted in MOLP medium, containing the appropriate antibiotics, and incubated it without shaking for 2 h at 30 °C. Subsequently, a portion of the bacteria diluted in MOLP was filled into specially manufactured glass capillaries with a thin tip of 10 μm in diameter. These capillaries were then placed with the large open end into the wells of the plate (the set-up is displayed in Figure 1b and Supplementary Figure 1). This incubation strategy created a stable, moist environment for the bacteria to form a biofilm within the very top of the capillary after 22-44 h of incubation time. The biofilm was also directly visible on the surface of the plate wells thanks to its characteristic pattern (Figure 1b), consisting of an assembly of wrinkles.

**Figure 1:**
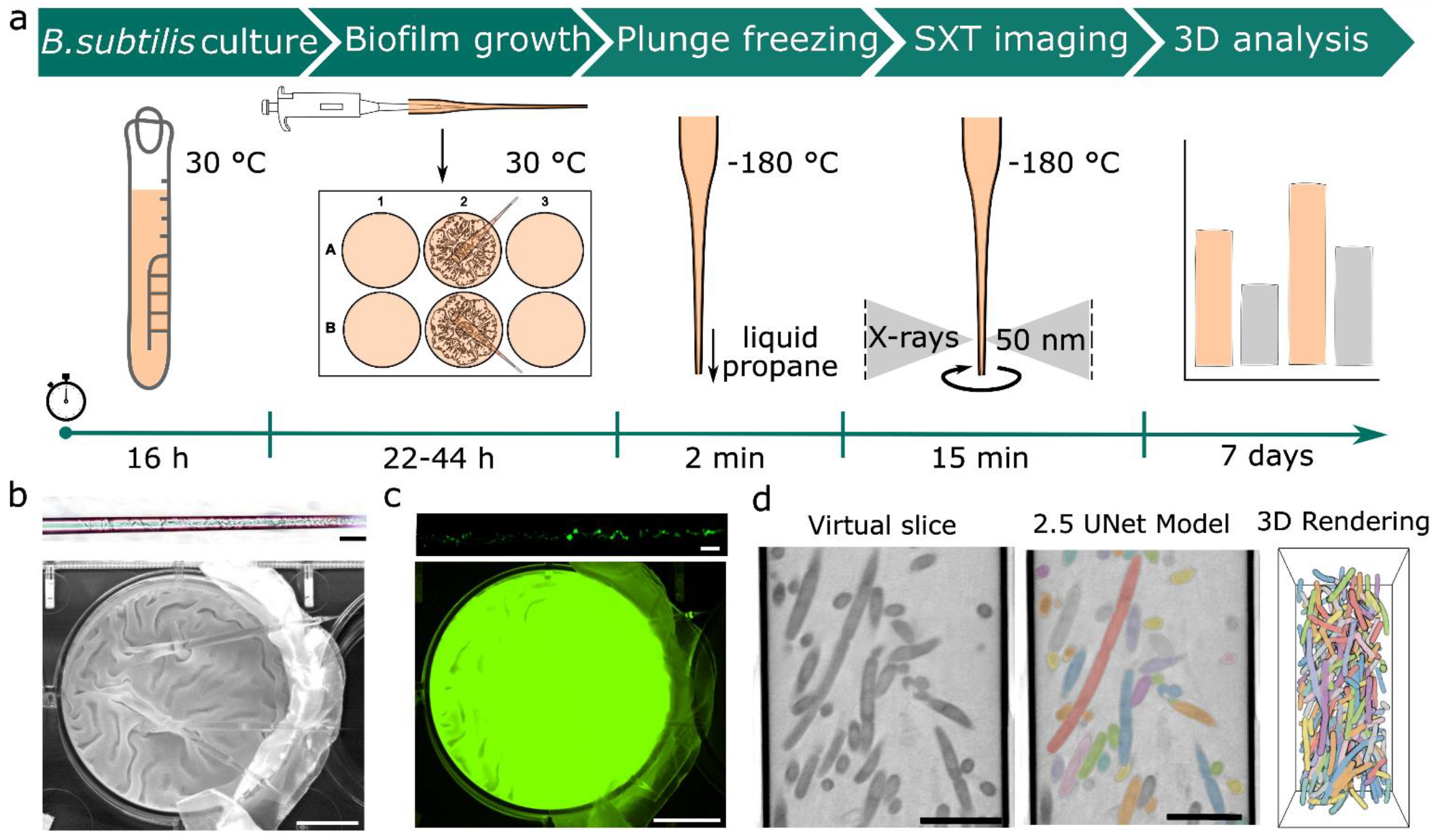
‘Biofilm-in-capillary’ workflow applied to GFP-expressing *B. subtilis*. **(a)** ‘Biofilm-in-capillary’ workflow steps are presented in a timeline. **(b)** Brightfield and **(c)** fluorescent images of GFP-expressing WT biofilm grown in the glass capillary (top image) and middle well (bottom image) of the 6-well plate setup after 22 h incubation. The folded structures made on the biofilm surface can also be shown inside the glass capillary. The scale bars on images from the well plate are 8.5 mm. The scale bars on images from the capillary are 40 μm. **(d)** For quantitative analysis of 3D morphometrical parameters of single bacteria and ECM fragments upon building biofilms, a neural network (2.5D UNet) supported by the Dragonfly software^26^ is trained on so-called virtual slices, which corresponds to a single plane of 32 nm thickness from the 3D SXT dataset. After automatic segmentation, the biofilm structure is segmented and rendered in 3D, as shown here with individual colours for each bacteria cell. Images are scaled at 5 μm.

The formation of biofilms within the capillaries was examined by light microscopy (Figures 1b top and 1c top), including *in vivo* imaging of GFP-expressing *B*.*subtilis*. Supplementary video 1 shows an example of individual bacterial cells migrating to the open tip of the capillary while other bacterial cells are stationary. Capillaries with detectable biofilm in the tip were cryo-preserved with a robotic plunge-freezer and imaged with the soft X-ray microscope as previously described^27^. The covered region of the tip is approximately 150-155 μm long. In this area, several series of SXT projection images are taken, from which 3D volumes containing the whole biofilm inside the capillary are calculated. The reconstructed volume was automatically segmented by the 2.5D U-Net neuronal network (Figure 1d), available as part of the Dragonfly software (version 2022.2)^26^. By segmenting the bacterial cells and the extracellular matrix (ECM), the biofilm structure was analyzed at both the single-cell level and gross morphology.

The established biofilm-in-capillary workflow enables the growth of bacterial biofilms from aerobe bacterial cells, compatible with high-resolution 3D imaging by SXT.

### TasA gene deletion leads to loss of cellular orientation and ECM compaction

At first, we applied the developed workflow to investigate wild-type *B. subtilis* biofilms. Such biofilms have a thickness of several hundreds of μm, with expected changes in their structure depending, for example, on oxygen concentration^28,29^. For this reason, we measured a considerable distance over the capillary length by collecting multiple fields of view (FOVs) (Figure 2a). The concomitant change of capillary diameter starting from the open tip was compensated for by gradually decreasing the magnification of the microscope and thus FOVs along the capillary length^23^. The acquired FOV close to the tip of the capillary had a starting dimension of 15μm x 15μm (see FOV 1 in Figures 2b and e). Continuing data collection along the length of the capillary for about 150-155 μm (Figures 2a and d), the FOV dimensions gradually increased from 15 μm to 17 μm for the last FOV (FOV 9 in Figure 2b). Examples of 2D virtual slices obtained from the reconstructed tomograms are shown in Figure 2b. The tomographic collection and reconstruction were then followed by segmentation and 3D rendering of individual bacteria and ECM, yielding the three-dimensional representation of the biofilm shown in Figure 2c (top). The individual features of the ECM are shown separately in Figure 2c (bottom).

**Figure 2:**
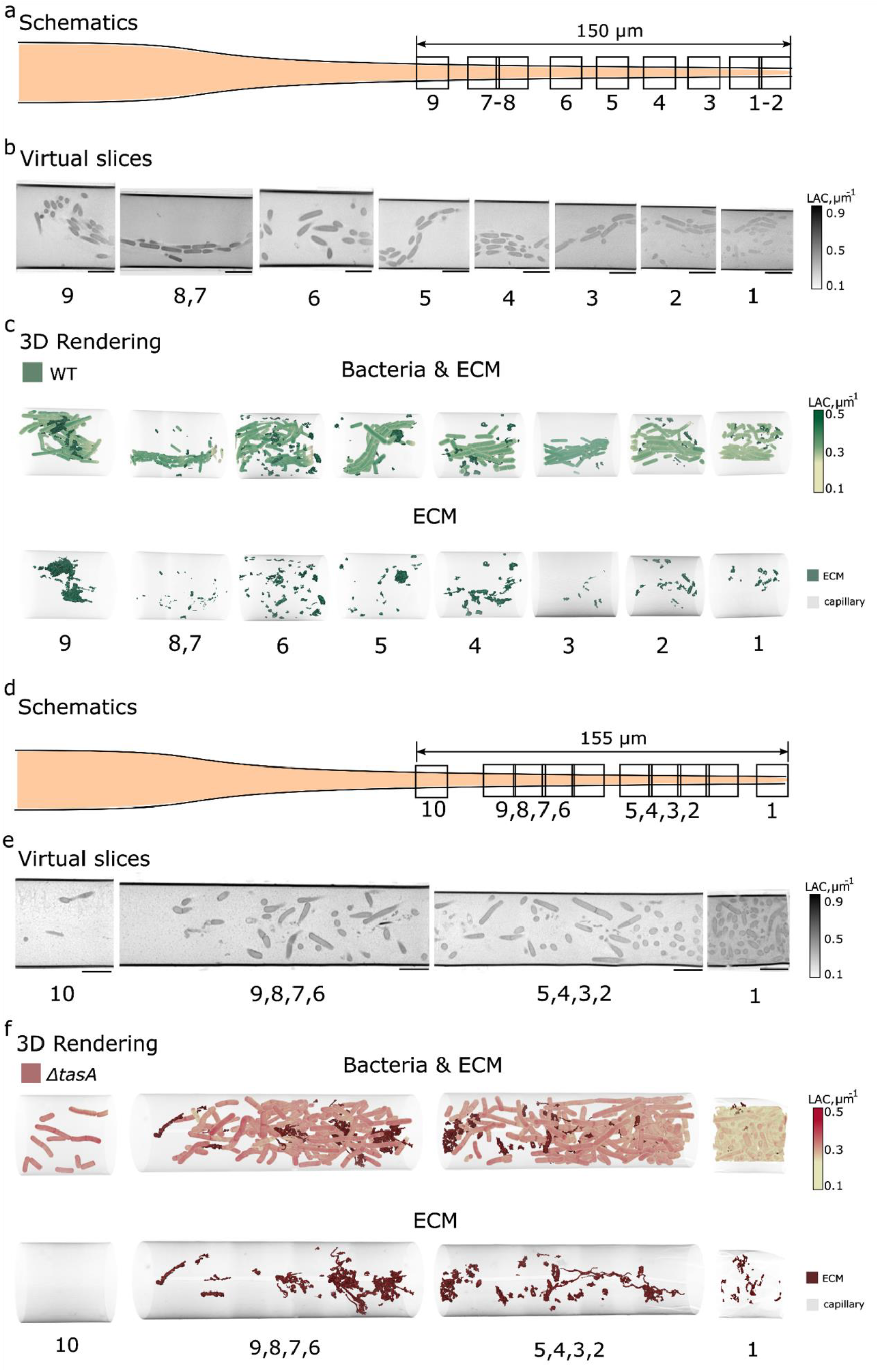
WT and Δ*tasA* bacteria built distinct biofilm structures. **(a)** Schematic of SXT imaging of WT *B. subtilis* biofilm (t=22 h). The biofilm extends up to 150 μm from the open capillary tip, as measured from the dimensions of the FOVs. **(b)** Two-dimensional virtual slices of FOVs 1-9 capture an oriented distribution of the bacteria along the glass capillary. Images are scaled at 5 μm. **(c)** 3D renderings of bacteria reveal the structured, helix-like, orientation of the bacteria along the capillary. The ECM fragments (dark green) show an uneven distribution of foam-like substructures along the capillary. **(d)** Schematic of SXT imaging of Δ*tasA B. subtilis* biofilm (t=22 h). The biofilm extends up to 155 μm from the open capillary tip of 10 μm width. **(e)** Two-dimensional virtual slices of FOVs 1-10 capture a loss of collective distribution of the bacteria along the glass capillary. Images are scaled at 5 μm. **(f)** 3D renderings of segmented bacteria reveal the loss of structured orientation of the Δ*tasA* bacteria along the capillary. The ECM fragments (dark red) show string-like structures that increase their volume along with the bacteria density.

Figure 2c (top) shows the bacteria clustering in tightly aligned patterns^30^, especially in FOVs 1,2,3, 4 and 8, with few ECM conglomerates inside the long, extended clusters (Figure 2c, bottom). The compactness and order of the bacterial alignment are striking. At the same time, FOVs 3,5,7, and 9 show chaining, apparent from turns of connected bacteria, see Supplementary Video 2 for a full 3D rotational view. These are the hallmarks of a biofilm in its early phase^31^, in agreement with the growth time of 22 hours. Figure 2c (bottom) shows the distribution of the ECM, which is more spread than in those FOVs that show intense clustering of bacteria.

The same procedure has been applied to cultures of *B*.*subtilis* with deleted *tasA* gene, termed Δ*tasA* (Figure 2d-f). In this case, it appears that the cells are no longer arranged in an organized chain fashion (Figure 2f top, Supplementary Video 3), they rather adopt random orientations. We conclude that TasA increases the level of biofilm organization. At the same time, larger areas filled with ECM are observed that sometimes show elongated, filament-like structures (Figure 2f bottom).

The above observations were followed up by a quantitative analysis at the cellular level. We compared the volumes, shapes (measured by aspect ratio), and density of individual bacterial cells. To ensure that *tasA* deletion is the sole factor that affects cellular phenotype in biofilms, we studied first *B*.*subtilis* WT and Δ*tasA* cells while supplying LB medium that does not support biofilm growth (Supplementary Figure 2), and found no volume difference. As time point t=0, the start of biofilm growth was chosen, assumed to happen 2 h after inoculation of the capillary and subsequent adaptation to the MOLP medium. This phenotype of single cells was then compared to the situation at t=22 h, see for both wild-type (Figure 3a) and Δ*tasA* (Figure 3b) cells. All measurements, that is bacteria’s volume, LAC, aspect ratio and ECM’s LAC are summarized in Supplementary Table 1.

**Figure 3:**
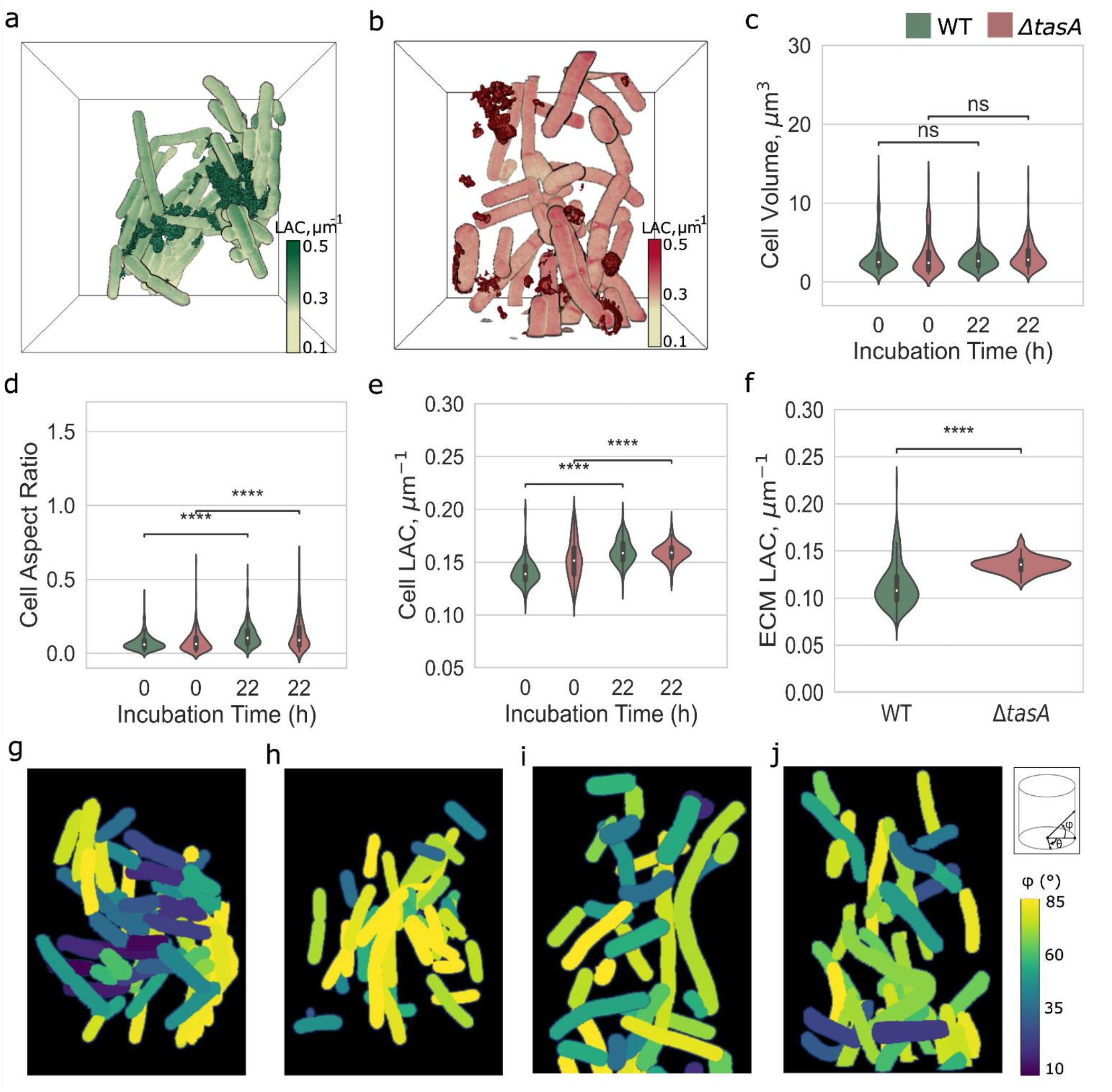
3D analysis of the WT and Δ*tasA* from single bacteria to grown biofilms. (**a**) 3D rendering of biofilms for WT (FOV 9; **Figure 2c**) and (**b**) ΔtasA (FOVs 2-5; **Figure 2f**) in a bounding box. The foam-like ECM (dark green) is wrapping the bacteria into a helix-like structure, while the thread-like ECM (dark red) is not attached to the bacteria and has no spatial orientation. The LAC is scaled from 0.1 μm^-1^ to 0.5 μm^-1^. Time-dependent comparison of cellular characteristics of single-cell bacteria (t=0 h) to biofilms (at t=22 h) for WT (green) and Δ*tasA* (red) genotypes is probed for (**c**) cell volume (μm^3^), (**d**) aspect ratio, and (**e**) LAC (μm^-1^). (**f**) X-ray LAC analysis of secreted ECM fragments. The bacteria orientation inside the capillary is measured by their orientation to θ ° and φ °, as sketched top right in panel (**j**). The φ ° shows orientation along the capillary and the θ ° is perpendicular to the capillary length. Here is presented the φ ° that bacteria acquire at WT (**g-h**) and Δ*tasA* (**i-j**) biofilm. Colors for φ ° are scaled from 10 ° to 85 °. Statistical analysis performed in t-test independent samples with Bonferroni correction, p-value (p) ns: 0.05 < p <= 1.00, (*): 0.01 < p <= 0.05, (^**^): 0.001 < p <= 0.01, (^***^): 0.0001 < p <= 0.001, (^****^): p <= 0.0001, with sample-size (n) for cell analysis, n=356 (WT, t=0 h), n=613 (Δ*tasA*, t=0 h), n=174 (WT, t=22 h) and n=274 (Δ*tasA*, t=22 h), respectively.

We observe that the average volume of *B*.*subtilis* cells is about 3 μm^3^ and does not change with *tasA* deletion, both in suspension and under biofilm growth conditions, see Figure 3c. While this average volume is not associated with the shape difference of bacteria grown in suspension, upon the formation of biofilm, cells become less elongated in comparison to cells grown in suspension condition, where the aspect ratio of cells increases from 0.07 ± 0.06 to 0.14 ± 0.10 in WT for t = 0 and t = 22, respectively; and from 0.10 ± 0.07 to 0.15 ± 0.14 for the same time points of a Δ*tasA* culture (Figure 3d, Supplementary Figure 2d). The differences in cell elongation as a consequence of defective cell division have been reported for cells in biofilms of *Pseudomonas aeruginosa*^32^. The changes in average volume and aspect ratio of cells are accompanied by a slightly increased LAC of WT cells from 0.14 ± 0.01 μm^-1^ to 0.16 ± 0.01 μm^-1^ for t = 0 and t = 22, respectively, whereas for Δ*tasA* (see Figure 3e and Supplementary Figure 2e) no significant difference was observed with corresponding values of 0.15 ± 0.02 μm^-1^ and 0.16 ± 0.01 μm^-1^. Such an increase in the X-ray absorption of the WT bacterial cells when building the biofilm indicates an increase in molecular density, which can be explained by metabolic changes. For example, enhanced protein and lipid oxidation have been previously observed for Δ*tasA* strains^33^. To understand whether the deletion of *tasA* contributes to the difference in composition of the secreted ECM, we compared the mean LAC of ECM fragments between the WT and Δ*tasA*. From the statistical analysis, we observe that the lack of the *tasA* gene results in chemically denser ECM fragments with a LAC of 0.14 ± 0.01 μm^-1^ as compared to 0.11 ± 0.02 μm^-1^ for WT. This 10 % density increase may be attributed, for example, to an increased number of carbon-rich proteins and sugars. Along these lines, the higher chemical density of ECM fragments in the case of Δ*tasA* might be caused by the elevated secretion of exopolysaccharides (EPS), due to cellular stress or increased reactive oxygen species (ROS) production of mutant cells^33^.

To analyze the organization of biofilms in more detail, we analyzed cell orientation along (φ°) and perpendicular (θ°) to the capillary directions (see Figure 3g-j and Supplementary Figure 3 for all FOVs). Assemblies of WT cells in biofilm show complex architectures along the tip of the capillary where in certain regions the neighbouring cells take identical orientation, see representative FOV in Figure 3g. The similarity in angular orientation together with chaining of bacteria results sometimes in a spiral geometry of biofilm. This suggests that during the growth of biofilms chains of bacteria search for orientation towards the oxygen-rich surface in a spiral, concerted manner as also visible in FOV 5 of Figure 2c, for example. These highly ordered regions are intercepted by regions of no order (Figure 3h), confirming the growth pattern of biofilms with varying ordered and disordered regions. On the contrary, in biofilms of the Δ*tasA* strain, all FOVs lack signs of ordered cell orientation, whether along or perpendicular to the capillary (Figure 3i-j, Supplementary Figure 3).

### Supplementing TasA protein to ΔtasA cultures restores biofilm morphology

Previous studies have shown that an external addition of TasA protein to cultures of Δ*tasA* bacteria restores the biofilm^5,34–36^ over 48 h at 30 °C^34^. We investigated this by applying our biofilm-in-capillary workflow over 44 h to understand the 3D organisation of such biofilms that are formed after addition of TasA protein. In the capillary setup, biofilms of Δ*tasA* cultures formed after addition of TasA show in principle the same morphology though not as pronounced as for WT *B*.*subtilis* (Supplementary Figure 4 in comparison to Figure 1b). In Fig. 4a, a representative FOV from SXT imaging of the Δ*tasA* culture shows a random arrangement of cells. The corresponding culture with TasA added (Fig. 4b) shows a compact cellular assembly, likely supported by the extended matrix. This packing of cells is similar to the wild-type biofilm but different to the Δ*tasA* situation shown in Fig. 4a. An extended chaining as displayed for WT biofilms in Fig. 2c is however not observed in the various FOVs of rescued biofilm.

**Figure 4:**
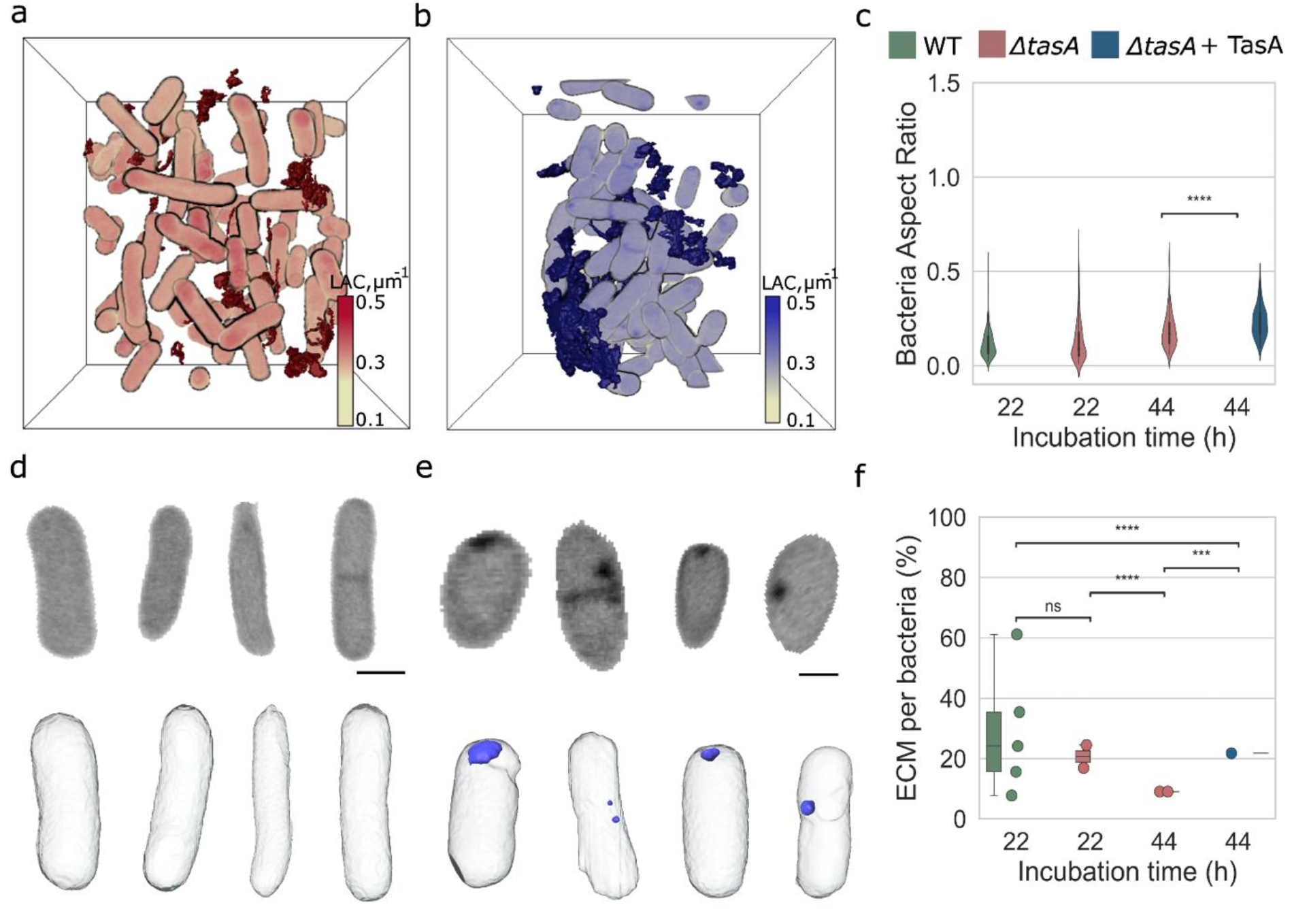
Restoration of biofilm upon extracellular addition of TasA to Δ*tasA*. **(a-b)** 3D rendering of the Δ*tasA* and Δ*tasA* treated with TasA (t=44 h) biofilm fragment in a bounding box. The ECM (dark red) fragments keep the same string-like structure as Δ*tasA* (t=22 h) biofilm, whereas the ECM fragments (dark blue) of Δ*tasA* cultures treated with TasA become denser. The LAC is scaled from 0.1 μm^-1^ to 0.5 μm^-1^. **(c)** Bacteria shape becomes significantly rounder upon TasA addition in Δ*tasA* (t=44 h). **(d-e)** The shape change is also visible in virtual slices and 3D renderings of individual bacteria from Δ*tasA* and Δ*tasA* + TasA cultures. Subcellular round and dense structures, here named “punctae” (light blue), are present in every treated with TasA bacterium. The scale bar is 1 μm. Virtual slice grayscale in LAC is from 0.1 μm^-1^ to 0.5 μm^-1^. **(f)** The extracellular addition of TasA restores by 13 % the ECM volume per number of bacterial cells as measured through SXT modality). Every dot represents each FOV considered for this analysis. Statistical analysis performed in t-test independent samples with Bonferroni correction, p-value (p) ns: 0.05 < p <= 1.00, (^*^): 0.01 < p <= 0.05, (^**^): 0.001 < p <= 0.01, (^***^): 0.0001 < p <= 0.001, (^****^): p <= 0.0001, with sample-size (n) for cell analysis, n=139 (WT, t=22 h), n=274 (Δ*tasA*, t=22 h), n=140 (Δ*tasA*, t=44 h) and n=103 (Δ*tasA* + TasA, t=44 h), respectively.

From the quantitative analysis (see also Supplementary Figure 5), the most striking effect at the cellular level is the change in cell shape (see Figure 4d and e). The Δ*tasA* cells treated with TasA acquire a spherical shape, measured by an increase of aspect ratio from 0.20 ± 0.01 in Δ*tasA* cultures to 0.24 ± 0.09 in biofilms of Δ*tasA* supplemented with TasA (see Figure 4c). This phenotype differs from the elongated WT cells (see Figure 3c). The biochemical density of the Δ*tasA* cells without supplement and in the biofilm occurring upon supplementation with TasA has significantly decreased to 0.15 ± 0.01 μm^-1^ from 0.16 ± 0.01 μm^-1^, respectively (see Supplementary Figure 5c). Surprisingly, most cells in the cultures containing TasA contain dense spherical structures, that we call “punctae”, with an LAC of 0.60 μm^-1^, see Figure 4e. While we have no precise information on the content and origin of these “punctae”, the LAC values suggest that the puncta contain a high amount of lipids, based on reported LAC for organelles in bacterial cells^37,38^.

To evaluate the level of ECM restoration by extracellular TasA, we measured the biochemical density of ECM fragments, see Figure 4f. In Δ*tasA* cultures treated with TasA, the ECM density (0.11 ± 0.01 μm^-1^) shows no difference in comparison to cultures containing untreated Δ*tasA* cells (0.10 ± 0.01 μm^-1^). Altogether, our data shows that the addition of extracellular TasA does not restore the phenotype of bacterial cells in biofilms including the ordered structure of cells in the biofilm. However, adding TasA to the growth media of the cells partially restores the composition of ECM.

## DISCUSSION

A “biofilm-in-capillary” workflow enables analysis of bacterial biofilms and quantitative measurements on both the subcellular and macroscopic size scales in a physiological state using full-rotation soft X-ray tomography. The workflow was validated on GFP-expressing *B. subtilis* biofilms for visualization of structural phenotypes and biochemical density of single cells and ECM within biofilms formed. The bacterium *Bacillus subtilis* is one of the best-studied model organisms for investigating biofilm formation. In addition to exopolysaccharides, *B. subtilis* biofilms contain TasA as major proteinaceous matrix component which is required for matrix formation, protection against oxidative stress, interaction with the membrane and it can also act as a developmental signal stimulating a subset of biofilm cells to revert to a motile phenotype^33,39^.

We, therefore, have focused on 3D imaging of WT *B. subtilis* biofilm and compared it to cultures of Δ*tasA* cells, restoring also the complex architecture of biofilms by addition of TasA protein. By use of full-rotation tomography and automatic segmentation based on machine learning, we were able to detect subtle differences in cellular phenotypes and ECM at statistically significant sample sizes. We could show that deletion of the *tasA* gene affects bacterial cells grown in suspension differently in comparison to the same bacteria during biofilm formation upon TasA supplementation. We observed significant changes in cellular elongation and chemical density as measured by soft X-ray absorption when *B. subtilis* cells are grown in biofilms. At the macroscopic level deletion of *tasA* leads to loss of cellular orientation. The extracellular addition of TasA protein to Δ*tasA* cultures partially restores the biochemical density and volume of the ECM; on the other hand, the cell organization and their phenotype are not rescued, suggesting complex functions in cellular metabolism and motility upon biofilm formation. Interestingly, based on high-spatial-resolution and sensitivity to chemical density, we were able to detect accumulation of lipids within Δ*tasA* cells upon TasA supplementation.

While we showed that biofilm-in-capillary workflow enables imaging of *B. subtilis* biofilms at high-spatial resolutions, it would be interesting to apply our approach to other types of biofilms, including those consisting of multiple species^40^, and interactions with artificial additives to biofilms^41^, for example, nanoparticles^42^. A potential extension of the proposed workflow would be localization of specific proteins by correlative fluorescence microscopy. Such correlation of structural information provided by soft X-ray tomography with functional aspects of specific proteins visualized via fluorescence microscopy has been demonstrated by several groups^43–46^.

Overall, due to high sensitivity to biochemical content SXT imaging combined with biofilm-in-capillary workflow enables unprecedented visualization and quantitative analysis of cells and ECM distribution within a 3D volume of biofilms. The combination of subcellular resolution and density measurements in 3D will provide better insights into the biofilm formation and microenvironment.

## METHODS

### Bacterial strain and culture conditions

The wild type and Δ*tasA* bacteria derive from the *B*.*subtilis* DK1042 strain based on pBS32-containing wild type (ancestral 3610 strain), which has an inducible translational fusion of ComI with Q-to-L change at codon 12 ^47^. For the integration of GFP-mut2 into the amyE locus under the control of Phyperspank, the gene and backbone (pDR111) were amplified by PCR. The plasmid was constructed using HIFI assembly (NEB #E2621). Transformations into *B. subtilis* wildtype (DK1042) and *ΔtasA* (tasA::kan) was performed as described previously^7^. Positive clones were selected on 150 μg/ml spectinomycin, and integration validated by PCR and sequencing.

Primers:

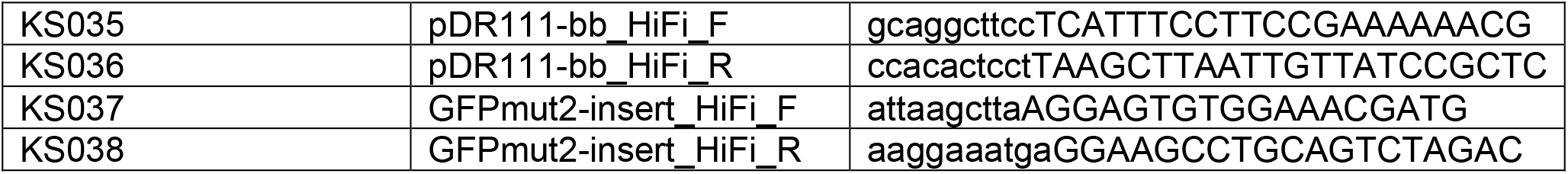

Bacteria cultures were grown from frozen stocks on Luria-Bertani (LB: 1 % tryptone, 0.5 % yeast extract, and 1 % NaCl) at 30 °C. After inoculation, bacteria were cultured overnight in LB medium at 30 °C under shaking at 160 rpm. Before performing biofilm assays, bacteria were subcultured in 1:100 concentration and left to cultivate to 1.0-1.2 optical density (OD) for 4-5 h. Biofilm formation was promoted by bacteria growth in the Medium optimal for lipopeptide production (MOLP)^25^: 30 g/L peptone, 20 g/L saccharose, 7 g/L yeast extract, 1.9 g/L KH_2_PO_4_, 0.001 mg/L CuSO_4_, 0.005 mg/L FeCl_3_.6H_2_O, 0.004 mg/L Na_2_MoO_4_, 0.002 mg/L KI, 3.6 mg/L MnSO_4_.H_2_O, 0.45 g/L MgSO_4_, 0.14 mg/L ZnSO_4_.7H_2_O, 0.01 mg/L H_3_BO_3_, 10 mg/L C_6_H_8_O_7_, adjusted to pH 7.0, for 22-44 h at 30 °C^25^. The final concentrations of antibiotics for *B*.*subtilis* bacteria, grown in either LB or MOLP medium, were 150 μg/mL spectinomycin (WT) and 10 μg/mL kanamycin (both *ΔtasA*). To inspect biofilm growth success inside the glass capillaries, bacteria were induced with 1 mM Isopropyl β-d-1-thiogalactopyranoside (IPTG) added to the MOLP medium to produce GFP.

### Single-cell-in-capillary workflow

Before seeding the wild type and Δ*tasA* bacteria for biofilm assay, we collected approximately 2*10^6^ bacteria/mL of each culture in an Eppendorf tube, which was spun down at 100x g for 3 min and resuspended in 20 μL of LB medium. The bacteria suspension was injected into thin-walled glass capillaries with an open tip of 9-10 μm width and cryopreserved as described below.

### Biofilm-in-capillary growth workflow

To promote biofilm formation into glass capillaries (32×1.5×1.05 mm, OD 1 ± 0.05, Hilgenberg GmbH), we prepared the following setup (Supplementary Fig 1): a 6-well plate with the middle cavity walls wrapped in decontaminated parafilm and two spacers taped at the side of the plate. When the bacterial culture reached the optimal cell density, the wild type and Δ*tasA* were seeded in the 6-well plate with 1:100 concentration in the MOLP, supplemented with antibiotics and IPTG. For the rescue experiment, TasA^28-261^ protein in 20 mM PO_4_ buffer pH 7,0 with 150 mM NaCl, was added to a final concentration of 200 μg/ml (8 μM) to MOLP. Bacteria were allowed to cultivate in biofilm formation setup for 2 h at 30 °C, and then, 20 μL of the bacteria-in-MOLP suspension was injected into 6 glass capillaries, with an open tip of 9-10 μm, per condition. Each capillary, after being filled up with bacteria in MOLP, was supported to stand by middle cavity walls wrapped with parafilm, and the spacers kept a safe distance between the capillary tips and the plate lid. To sustain the humidity levels desirable for the *B*.*subtilis* biofilm formation, we wrapped the 6-well plate with wet towels in a sealable plastic bag and incubated for 22-44 h at 30 °C.

In the course of the rescue experiments, it was noted that different recombinant TasA preparations support biofilm formation to a different extent. Older preparations, frozen or stored at higher concentrations support biofilm formation better than freshly prepared monomeric TasA.

### Light microscopy

Before proceeding with cryo-soft X-ray tomography, we inspected the quality of the plates and capillaries containing biofilms. For the inspection of the biofilm grown into the 6-well plates, we imaged the plates at a Fusion FX device (Vilber) with 120 ms exposure time. To examine the biofilm growth into the capillaries, we acquired a time-lapse of bacteria forming biofilm close to the tip, at 320 s exposure time for 500 ms with a 488 nm laser and a 20x objective, by an inverted (Carl Zeiss AxioVision CD25) epi-fluorescence microscope. The *in vivo* imaging occurred in the National Laboratory for X-ray Tomography (NCXT) at Lawrence Berkeley National Laboratory, USA.

### Cryo soft X-ray tomography

The capillaries containing biofilms were rapidly frozen by immersion in liquid propane cooled with liquid nitrogen (∼ -90 °C). Soft X-ray tomographic data were acquired through full-rotation imaging using the soft X-ray microscope, XM-2, at the National Center for X-ray Tomography, housed at the Advanced Light Source of Lawrence Berkeley National Laboratory in Berkeley, CA (https://ncxt.org/). To prevent radiation damage, cells were exposed to a stream of liquid-nitrogen-cooled helium gas during the data collection process. Each dataset involved the collection of 184 ° rotation tomographs^48^, with one projection image captured per 2 ° angle^19,27^. The automatic reconstruction software was employed for the automatic reconstruction of projection images into 3D volume^49^. This process combined the information from 92 slices over 184 ° around the capillary, followed by segmentation and 3D rendering, to visualize bacteria and biofilm in 3D.

### Image Analysis

The light microscopy images of biofilms grown into plates and capillaries were analyzed with the open-source Fiji software^50^. For the automatic segmentation of single bacteria, the reconstructed soft X-ray tomography datasets were submitted to the 2.5 UNet model, supported by DragonFly (version 2022.2)^26^. The ECM was segmented manually based on the pixel intensity threshold with Amira 2020.3.1 software. The 3D renderings of segmented single bacteria, ECM, and capillary walls were prepared with Amira. Bacteria and ECM detected close to the tip (FOV 1 in Figure 2b and 2e) are excluded from the 3D quantitative analysis because this area is compacted with bacteria and there is a lack of contrast important for the ECM precise segmentation.

### Statistical Analysis

Statistical analysis and plots on 3D analyzed soft X-ray tomography datasets were prepared with Jupyter notebook, using the statannot package (https://github.com/webermarcolivier/statannot) to compute statistical tests (Student’s t-test) and add statistical annotations^51^. Figures were assembled with Inkscape 1.3 software.

## DATA AVAILABILITY

The data is available from the corresponding author upon request.

## ACKNOWLEDGEMENTS

V.W.’s work was funded within the framework of the Excellence Strategy of the Federal and State Governments of Germany, the Excellence Cluster “3D Matter Made to Order” (3DMM2O), the CLEXM MSCA-DN project funded by the European Union under Horizon Europe and by the CoCID project (no. 101017116) funded within EU Research and Innovation Act. *A*.*C*. conducted the experiments at NCXT, by being supported by *the Joachim Herz Stiftung Foundation’s program for ‘Add-on Fellowships for Interdisciplinary Life Science’, cohort 2022*. C.L. and the NCXT were supported by NIH NIGMS P30GM138441 and DOE Biological and Environmental Research Project DE-AC02-05CH11231.

## AUTHOR CONTRIBUTIONS

V.W. and A.D. conceived the presented idea. A.C., B.V., and A.D. performed the development of the “Biofilm-in-capillary” growth protocol. K.D. and A.D. developed bacteria strains. A.C., J-H.., V.L. performed SXT experiments. A.C., A.T.H. and K.K. performed segmentation and statistical analysis. M.A.L.G., C.L., K.T., H.O. and V.W. supervised the fundings of this work. All authors discussed the results and contributed to the final manuscript.

## COMPETING INTERESTS

There is no conflict of interest.

## ADDITIONAL INFORMATION

## Notes

### Competing Interest Statement

The authors have declared no competing interest.

